# Cycling and prostate cancer risk – Bayesian insights from observational study data

**DOI:** 10.1101/2020.06.25.171546

**Authors:** Benjamin T. Vincent

## Abstract

Men have a very high lifetime risk of developing prostate cancer, and so there is a pressing need to understand factors that influence this risk. One factor of interest is whether cycling increases of decreases prostate cancer lifetime risk. Two large observational studies of cyclists noted very low rates of prostate cancer amongst cyclists relative to the general population – neither however drew causal conclusions about risk based on this observational prevalence data alone. Here we explore if and how we can use such data to update our beliefs about whether cycling increases or decreases prostate cancer risk – we use probabilistic methods to quantify belief in risk given the observational data available. We examine whether there is a dose– response relationship, how we can make inferences about risks, and the impact upon selection bias upon these inferences. A simple analysis leads us to believe that cycling decreases risk, but we show how this is mistaken unless selection bias can be ruled out. If cyclists who develop prostate cancer are less likely to respond to these surveys, we may be mislead into believing that cycling decreases risk even if it actually increases risk. Overall we explore precisely why it is hard to draw conclusions about risk factors based upon observational prevalence data.

## Introduction

We discuss two large, cross-sectional, observational, surveys of male cyclists. The first by Hollingworth, Harper, and Hamer (2014) reports results deriving from the 2012-13 Cycling for Health UK study which consisted of 5,282 male cyclists. As detailed below, this study claimed a dose-response relationship between amount of cycling per week and prostate cancer risk. The second study by Koupparis et al. (2020) achieved a sample of 8,074 male cyclists with the aid of the Global Cycling Network (GCN). This study claimed cycling does not influence prostate cancer risk.

Because of the high lifetime prevalence of prostate cancer (17.9% in the UK, Cancer Research UK, 2020a) and the popularity of cycling, these studies have attracted significant attention – the former was covered by UK national press and the latter was covered by GCN in a video https://www.youtube.com/watch?v=Nh2WraVt4JE. This video (which has been viewed approximately 89,000 times at time of writing) sends a potentially overconfident message about cycling and prostate cancer risk. While neither study claims proof of a causal link (in either direction), the medical and public interest is such that we believe the notably low prostate cancer rates in respondents relative to the general population deserves further investigation. Using a Bayesian probabilistic perspective, we explore: a) if we should believe in a protective effect, b) how selection bias may interfere with data driven beliefs about risk, and c) if we should believe in a dose-response relationship between cycling volume and risk.

## Protective effect?

Consider the Bayesian Network in Figure 1 which depicts a possible influence of cycling upon prostate cancer. In order to understand whether cycling is a risk factor for prostate cancer we can calculate the relative risk, defined as

**Figure 1:**
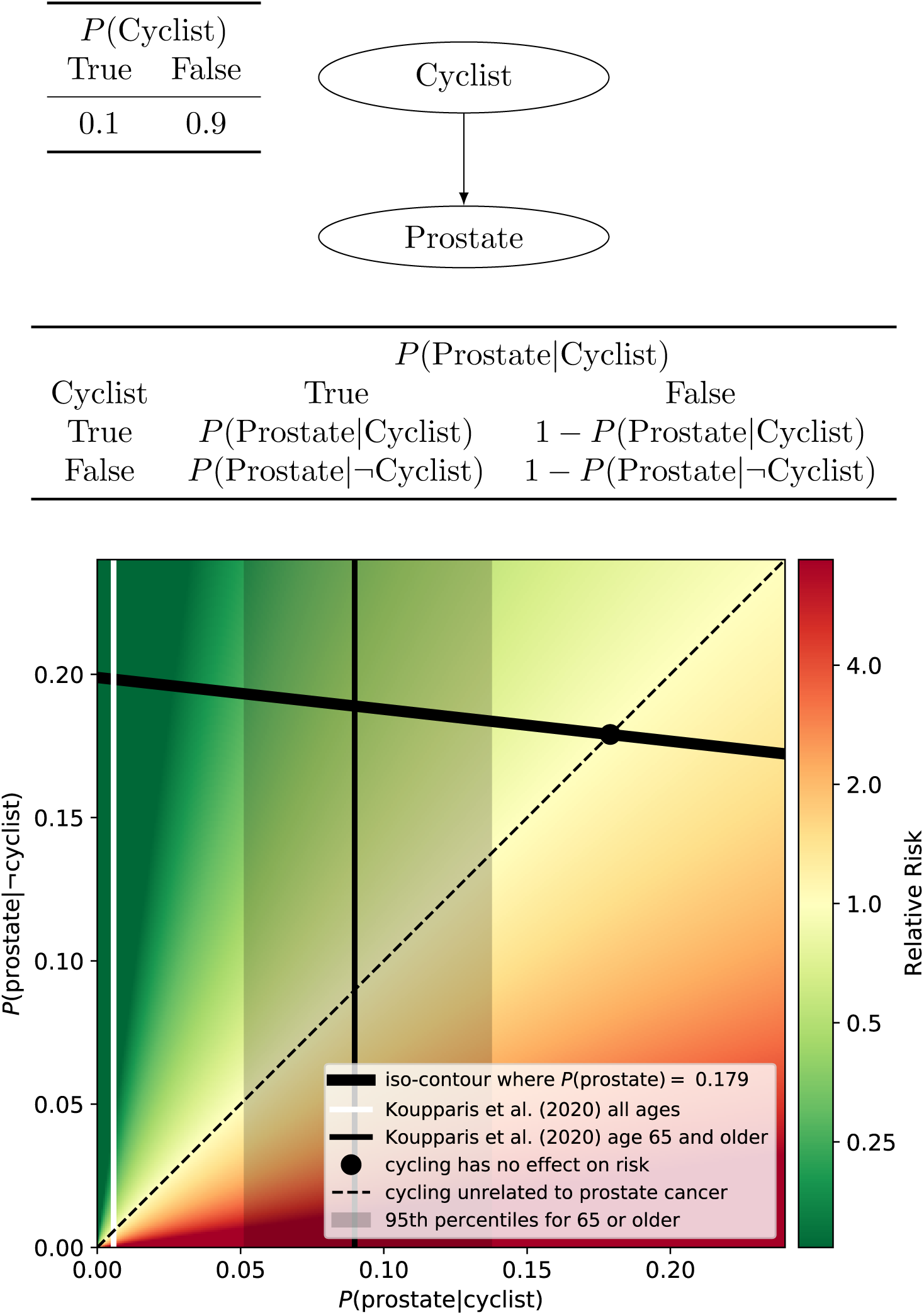
A simple Bayesian Network (top) where cycling may influence the probability of prostate cancer. The bottom plot shows a range of sets of probabilities of getting prostate cancer given one is a cyclist or not a cyclist. Probabilities shown by the thick black line are consistent with lifetime risk of prostate cancer (17.9%). Regions above the diagonal correspond to cycling increasing prostate cancer risk and regions below the diagonal correspond to cycling decreasing prostate cancer risk. Rate of prostate cancer in cyclists in the upper age category of Koupparis et al. (2020) is shown by the solid vertical line, with 95% credible intervals indicated by the dark shaded region.

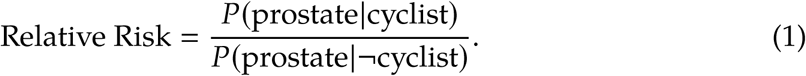

That is, we need to know both the probability of having prostate cancer amongst cyclists, *P* (prostate|cyclist), and the probability of having prostate cancer in non-cyclists, *P* (prostate|¬cyclist). We can obtain estimates of *P* (prostate|cyclist) from observational studies measuring the frequency of prostate cancer in cyclists, such as Hollingworth et al. (2014) and Koupparis et al. (2020). Without direct measurement of the risk of prostate cancer in non-cyclists, we can estimate this by first noting that the overall probability of having prostate cancer, *P*(prostate) is defined as

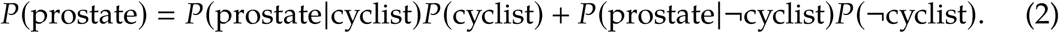

and then rearrange this equation to result in

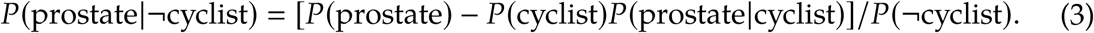

We can now gather estimates for the terms on the right hand side of the equation in order to estimate *P* (prostate|¬cyclist) and thus estimate the relative risk. First, it is not easy to precisely estimate the lifetime proportion of men who can be considered as cyclists – however based on UK Government statistics, we make the reasonable approximation of 10% (Department for Transport, 2018), that is, *P* (cyclist) = 0.1, and *P* (¬cyclist) = 0.9. Second, we can take the lifetime risk of prostate cancer in the UK as 17.9%, that is *P* (prostate) = 0.179 (Cancer Research UK, 2020a). Third, we can take the probability of prostate cancer in cyclists from Koupparis et al. (2020) as 0.57%, that is *P* (prostate|cyclist) = 0.0057.

Figure 1 visualises all of this information – the thick black line represents a set of combinations of risks which are consistent with the overall lifetime risk (see Equation 3). Given our initial estimate of *P* (prostate|cyclist) = 0.0057 from Koupparis et al. (2020) (white line in Figure 1), then we can calculate *P* (prostate|¬cyclist) = 0.198 using Equation 3, and thus estimate the relative risk as an 0.0057/0.198 = 0.029. This equates to an astonishing 34.8 fold *decrease* in risk of prostate cancer in cyclists. But is this valid?

It is clear, due to the wide age distribution of participants in the survey, that the 0.0057 estimate is a lower bound and is not equal to the lifetime risk. That is, some of the participants may well go on to acquire a prostate cancer diagnosis in later life. So it would be more appropriate to compare prostate cancer rates of cyclists and non-cyclists, in age-matched categories. The survey of Koupparis et al. (2020) only targeted cyclists and so the corresponding rates for non-cyclists are not available – so as a reasonable approximation we could take the rate of prostate cancer in cyclists in the highest age bracket (age 65 and above) as an estimate of *P* (prostate|cyclist) more comparable to the lifetime risk. This results in *P* (prostate|cyclist) = 0.098 (9.8%, taken from their Figure 4a where 15 of 167 participants report prostate cancer diagnoses). Inserting into Equation 3 results in an estimate of *P* (prostate|¬cyclist) = 0.188 (18.8%, thin black line in Figure 1). This results in a relative risk of 0.098/0.188 = 0.521, that is cycling may reduce risk of prostate cancer. There are a few caveats however.

The first is that there were only 167 respondents age 65 or over, so it is important to take our uncertainty of *P* (prostate|cyclist) into account – we can describe our beliefs as *P* (prostate|cyclist) ∼ Beta (15, 152). This results in 95% Bayesian credible intervals of 0.051 – 0.137 and is shown by the shaded region in Figure 1. We can see that the black point (representing cycling being unrelated to lifetime prostate cancer risk) is outside of our 95% credible range of beliefs. That is, based on reasonable assumptions, the relative risk is 0.475 [*CI*_95%_ : 0.266, 0.749] ^1^, and so we still have provisional evidence for a protective effect of cycling.

The second is that the calculations above do not represent all of our uncertainties. For example we have taken *P* (cyclist) = 0.1, but in reality we do have uncertainty about what this probability is. We also have uncertainty over the true probability of a cyclist having prostate cancer, *P* (prostate|cyclist). We can obtain improved estimates in the relative risk by taking into account all of our uncertainties using Monte Carlo methods. We show (in the Supplementary Material) that even when taking these additional uncertainties into account we retain our provisional belief that cycling decreases prostate cancer risk.

The third is that the higher estimate of *P* (prostate|cyclist) = 0.098, from participants age 65 and older, is potentially still an underestimate. The highest rates of prostate cancer are in the 75-79 age group and remains high in older age categories (Cancer Research UK, 2020b), and it is not clear how many of these older participants are reflected in the survey.

In summary, one way to understand the notably low rate of prostate cancer in the Koupparis et al. (2020) survey relative to the lifetime risk, is to recognise that the former is ‘diluted’ by younger cyclists who may go on to develop prostate cancer in older age. Using data from cyclists in the highest age category in the study is more comparable to the lifetime risk figure. Doing this however, still provides provisional justification to believe that cycling offers a protective effect. However, the next section explores one possible reason why lower rates of prostate cancer risk in cyclists may tell us very little (if anything) about cycling and prostate cancer risk.

## Selection bias?

One of the key characteristics of the datasets from Hollingworth et al. (2014) and Koupparis et al. (2020) is that participants were cyclists. So in interpreting the very low rate of prostate cancer in survey respondents, we must also take possible selection bias into account. We can add this possibility into our understanding by extending the Bayesian Network in Figure 1 to include whether or not a participant is included in the survey (see Figure 2). The conditional probability table governing the probability of being in the survey given cycling and prostate status in the figure embodies the hypotheses that: a) of all cyclists, there is a relatively low probability of completing the survey^2^, but non-cyclists are not included in the survey, and b) cyclists with prostate cancer are 4 times less likely to be included in the survey than cyclists without prostate cancer^3^.

**Figure 2:**
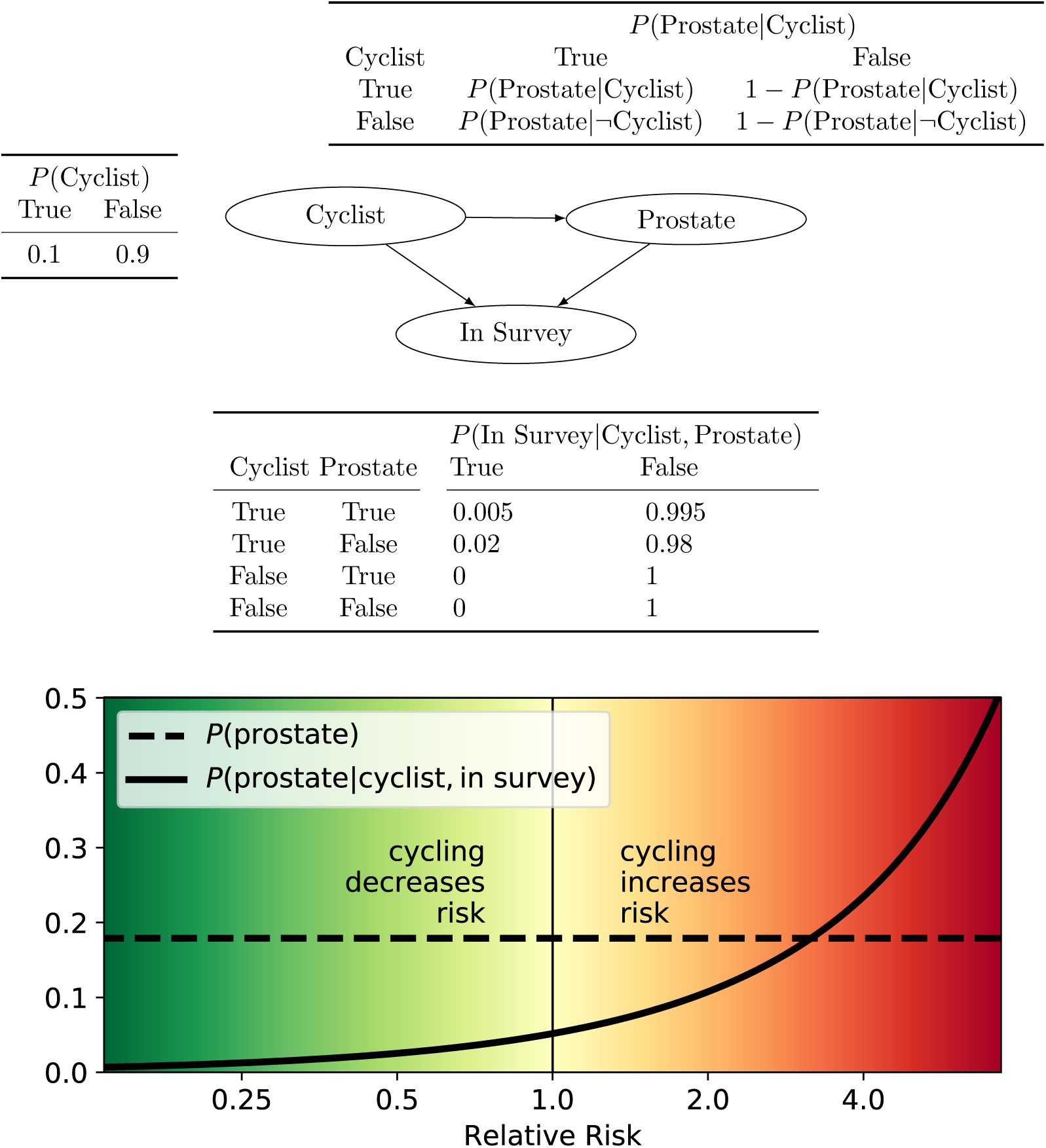
The effects of selection bias. (Top) A Bayesian network where cycling may have some influence upon prostate cancer risk (as in Figure 1), and inclusion in the survey is influenced by both cycling and prostate cancer status. (Bottom) We can see the effect of selection bias is that risk of prostate cancer in cyclists in the survey, *P* (prostate|cyclist), in survey, will be lower than the population lifetime risk, *P* (prostate), even if cycling has a positive relative risk (up to about a relative risk of ∼3.5). Calculated using the bnlearn (Taskesen, 2019) and pgmpy (Ankan & Panda, 2015) toolboxes.

This latter hypothesis is they key one of interest – we must take seriously the possibility that cyclists with prostate cancer are less likely to complete the survey than those without prostate cancer. One reason for this could be that cyclists who develop prostate cancer stop cycling (of their own volition or because of medical advice) and are less likely to take part in surveys aimed at cyclists. Another could be because of psychological avoidance of the uncomfortable medical situation they face. Regardless, I do not attempt to prove that this hypothesis is correct, I simply show that *if* it is the case, then we are likely to find low rates of prostate cancer in survey respondents regardless of whether cycling is protective, irrelevant, or a risk factor for prostate cancer.

Figure 2 (bottom) shows the probability of a cyclist included in the survey will have a prostate cancer diagnosis as a function of level of risk. This is plotted alongside the baseline prevalence in the population. We can see, somewhat unintuitively, that we should expect a very low proportion of cyclists in the survey to have prostate cancer relative to the population baseline. And that is true when cycling decreases risk, plays no role in risk (vertical line at zero), or increases risk of prostate cancer. The exception is when cycling represents a *very* large relative risk (over approximately 3.5) then the proportion of cyclists in the survey with prostate cancer will be higher than in the general population.

In summary, *if* there is a chance that cyclists (or ex-cyclists) with prostate cancer have a lower rate of survey completion than cyclists without prostate cancer (for any reason), then we should expect lower rates of prostate cancer in survey respondents than the general population. It is therefore crucial for any claims that cycling increases, does not effect, or decreases prostate cancer risk based upon surveys such as these, take selection effects into account.

## Dose–response relationship?

Hollingworth et al. (2014) analysed a subset of (2,027) men over 50 in their sample and show a positive relationship between cycling rate (hours/week) and probability of occurrence. However, because this dataset only consisted of 36 men who declared a prostate cancer diagnosis we must ask how much confidence can we place on this apparent dose-response relationship?

Figure 3 shows our beliefs over the probabilities of prostate cancer for a given cycling rate, *P* (prostate|cycling rate), based upon the reported relative frequency of cases. This kind of Bayesian approach is useful and intuitive – we can assess how much we believe that rates are different amongst different groups based upon overlap of 95% credible intervals^4^(see for example Kruschke & Liddell, 2017). We can see that the 95% credible intervals are broad relative to the differences between groups – judging in this way would lead us to consider only a difference in prostate cancer prevalence between the lowest and the highest rate of cycling groups. And given that this lowest category only includes 3 cases, it might be imprudent to conclude in a definitive dose-response relationship. It is important to note that the 95% credible intervals in Figure 3 represent the range of credible intervals *if* we believe prostate diagnosis responses are 100% correct for all participants. Given the low numbers of respondents, even if we have a small degree of uncertainty over response accuracy then the 95% credible intervals would be much broader. Based on this kind of analysis, the data should *not* lead us to strongly believe in a dose-response relationship.

**Figure 3:**
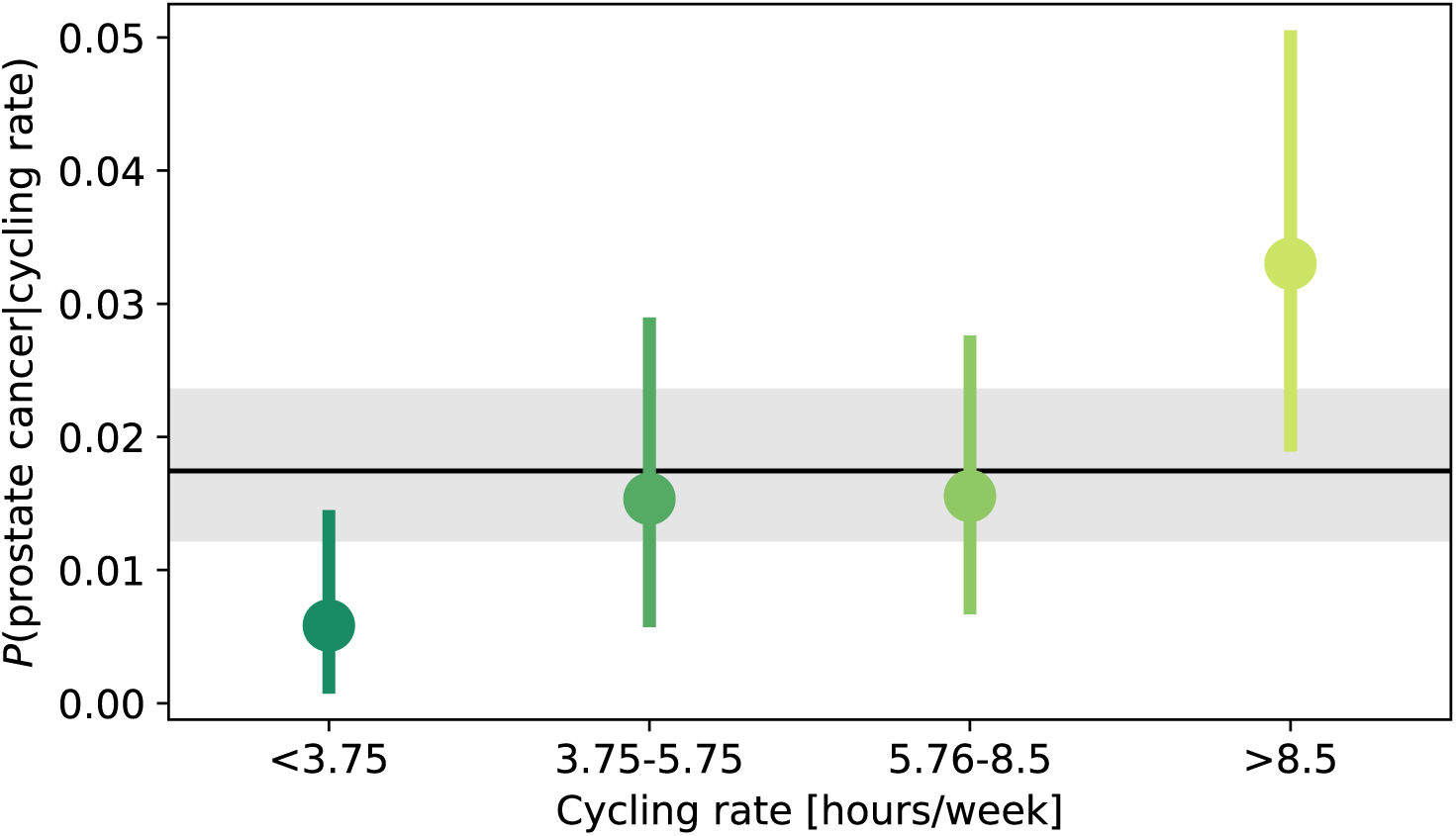
Probability of prostate cancer for a given cycling rate (hours/week). Points and lines represent the expectation and 95% Bayesian credible intervals. Black line and shaded region represent the expectation and 95% Bayesian credible interval for probability of prostate cancer overall. This plot is based solely upon frequency of cases in Hollingworth et al. (2014), using Beta distributions to represent our beliefs.

The logistic regression analyses by (Koupparis et al., 2020) can also address the question of whether there is a dose-response relationship. This kind of analysis informs us whether it is possible to predict prostate cancer diagnoses of cyclists based upon predictor variables such as years cycled, cycling frequency, hours cycled per week, and additional confounding variables such as age and Body Mass Index. They do not report any significant p-values for these analyses, which suggests no dose-response relationship. However, there are two reasons why we may wish to retain some uncertainty in our minds. Firstly, because of the very low rate of prostate cancer in the sample, we have an extreme asymmetry in the number of data points of those with no cancer (8,027) and those with prostate cancer (47). So an open question is whether these 47 respondents with prostate cancer are truly representative of cyclists with prostate cancer in general. Analyses which provide degrees of certainty, such as Bayesian or statistical boostrap methods (Efron & Tibshirani, 1986), would improve this situation. Second, because all respondents were cyclists, the dataset is subject to the same potential selection bias as discussed above. It *could* be that there is a dose-response relationship and those who go on to develop prostate cancer stop cycling (for whatever reason) and are therefore not included in these datasets. Before one could strongly conclude lack of a dose-response relationship, in the same way as the previous section, one would have to convincingly rule out any such selection bias. In summary, it seems prudent to conclude that while the data examined does not support a dose-response relationship, neither is there strong evidence to definitively conclude that there is no dose-response relationship.

## Conclusions

The findings from two large observational studies come to conflicting conclusions about the effect of cycling on prostate cancer risk. Hollingworth et al. (2014) propose a dose-response relationship between hours/week of cycling and risk of prostate cancer in men over 50. When we consider that the numbers of survey respondents with prostate cancer were low and the corresponding Bayesian 95% credible intervals, we conclude that it we do not yet have compelling evidence for a dose-response relationship. Based on logistic regression analyses, Koupparis et al. (2020) claim no dose-response relationship. But when we consider the number of respondents with prostate cancer was low, and the potential for selection bias, we find it hard to firmly conclude there is no dose-response relationship.

Both studies report very low prostate cancer risks for cyclists, much lower than the overall lifetime risk in the general population. It was shown that if cyclists (or ex-cyclists) with prostate cancer are less likely to take part in these surveys, then we we should expect to see lower prostate cancer risks in a cycling group as compared to the general population, even if cycling increases prostate cancer risk.

Overall, while we have two large observational studies which inform the debate, we conclude that it is too early to draw definitive conclusions either way about cycling and prostate cancer risk. It is important that this is considered when the media translate scientific studies for public consumption.

## Acknowledgements

The author thanks Kimberley Morrison for feedback on an earlier version of this manuscript.

## Appendix

All data, code, and Supplementary Materials are available from the Open Science Foundation, https://osf.io/fjahw/.

## Footnotes

1 Note that this estimate is based on the assumption that *P*(cyclist) = 0.1.

2 The exact probability cyclists with prostate cancer being included in the survey is not important. Rather, it is the relative probability of cyclists with and without prostate cancer being included in the survey which is poignant here.

3 The exact ratio used here will affect the curve in Figure 2, but the focus is on demonstrating the point, rather than making a claim about the exact value.

4 Bayesian 95% credible intervals define a range in which you are 95% sure contains the true value. This contrasts to the Frequentist 95% confidence intervals in which in 95% of theoretically repeated experiments to contain the true value (Morey, Hoekstra, Rouder, Lee, & Wagenmakers, 2016)

## References

Ankan, A., & Panda, A. (2015). pgmpy: Probabilistic graphical models using python. In Proceedings of the 14th python in science conference (scipy 2015).

Cancer Research UK. (2020a). Lifetime risk by cancer type and sex. https://www.cancerresearchuk.org/health-professional/cancer-statistics/risk/lifetime-risk#heading-One. (Accessed: 2020-06-17)

Cancer Research UK. (2020b). Prostate cancer incidence by age. https://www.cancerresearchuk.org/health-professional/cancer-statistics/statistics-by-cancer-type/prostate-cancer/incidence#heading-One. (Accessed: 2020-06-17)

Department for Transport. (2018). Walking and cycling statistics, England: 2018. https://www.gov.uk/government/statistics/walking-and-cycling-statistics-england-2018. (Accessed: 2020-06-17)

Efron, B., & Tibshirani, R. (1986). Bootstrap methods for standard errors, confidence intervals, and other measures of statistical accuracy. Statistical Science.

Hollingworth, M., Harper, A., & Hamer, M. (2014). An Observational Study of Erectile Dysfunction, Infertility, and Prostate Cancer in Regular Cyclists: Cycling for Health UK Study. Journal of Men’s Health, 11(2), 75–79.

Koupparis, A., Mehmi, A., Rava, M., Kearley, S., Aning, J., Rowe, E., & Richardson, S. (2020). Cycling and men’s health: A worldwide survey in association the Global Cycling Network. Journal of Clinical Urology, 161, 205141582091538–7.

Kruschke, J. K., & Liddell, T. M. (2017). Bayesian data analysis for newcomers. Psychonomic Bulletin & Review, 17(3), 251–23.

Morey, R. D., Hoekstra, R., Rouder, J. N., Lee, M. D., & Wagenmakers, E.-J. (2016). The fallacy of placing confidence in confidence intervals. Psychonomic Bulletin & Review, 23, 103–123.

Taskesen, E. (2019). bnlearn. https://github.com/erdogant/bnlearn.

